# CD8+ cells and small viral reservoirs facilitate post-ART control of SIV in Mauritian cynomolgus macaques

**DOI:** 10.1101/2023.03.01.530655

**Authors:** Olivia E. Harwood, Lea M. Matschke, Ryan V. Moriarty, Alexis J. Balgeman, Abigail J. Weaver, Amy L. Ellis-Connell, Andrea M. Weiler, Lee C. Winchester, Courtney V. Fletcher, Thomas C. Friedrich, Brandon F. Keele, David H. O’Connor, Jessica D. Lang, Matthew R. Reynolds, Shelby L. O’Connor

## Abstract

Sustainable HIV remission after antiretroviral therapy (ART) withdrawal, or post-treatment control (PTC), remains a top priority for HIV treatment. We observed surprising PTC in an MHC-haplomatched cohort of MHC-M3+ SIVmac239+ Mauritian cynomolgus macaques (MCMs) initiated on ART at two weeks post-infection (wpi). For six months after ART withdrawal, we observed undetectable or transient viremia in seven of eight MCMs. In vivo depletion of CD8α+ cells induced rebound in all animals, indicating the PTC was mediated, at least in part, by CD8α+ cells. We found that MCMs had smaller acute viral reservoirs than a cohort of identically infected rhesus macaques, a population that rarely develops PTC. The mechanisms by which unusually small viral reservoirs and CD8α+ cell-mediated virus suppression enable PTC can be investigated using this MHC-haplomatched MCM model. Further, defining the immunologic mechanisms that engender PTC in this model may identify therapeutic targets for inducing durable HIV remission in humans.

## Introduction

Identifying the immunological and virological attributes that lead to antiretroviral therapy (ART)-free HIV remission is critical for establishing a functional cure. A rare group of people infected with HIV can suppress virus replication to ≤400 copies/mL for ≥16 weeks to several years after ART interruption. These individuals are termed post-treatment controllers (PTCs)^1–4^. While specific major histocompatibility complex (MHC) alleles are associated with spontaneous control of HIV without ART (elite control; EC), MHC alleles are not thought to influence the likelihood of becoming a PTC^2,5–9^. The underlying mechanisms that lead to PTC are poorly understood. Unfortunately, it is challenging to study PTC in humans since people with HIV rarely undergo treatment interruption, PTCs are uncommon, and PTC cohorts are heterogeneous in genetic composition, timing of ART initiation, and definition of PTC^10^.

Among PTCs, the most common intervention is starting ART early after infection (reviewed in^11^). This phenomenon is best characterized in the VISCONTI, CHAMP, SPARTAC, and CASCADE cohorts, where 4-16% of participants became PTCs^1–4^. Unlike post-exposure prophylaxis, which can prevent establishment of systemic HIV infection if ART is initiated within 72 hours of exposure^12^, initiating ART within weeks of exposure limits the size of cell-associated HIV-1 DNA reservoirs^13,14^ and prevents the accumulation of viral variants, while preserving antiviral immune function^15,16^. However, not everyone who begins ART immediately after infection becomes a PTC; it is unclear why early ART potentiates PTC in some individuals and not others.

Because CD8+ T cells mediate viral load decline after peak viremia both on and off ART^17,18^ and suppress viral replication during ART^19^, CD8+ T cells may also contribute to PTC. Unlike the well-established role of CD8+ T cell-mediated virus suppression observed in most ECs^20,21^, PTCs generally have low numbers of IFN-γ-producing HIV-specific CD8+ T cells that poorly suppress HIV replication ex vivo^2^. While T cell exhaustion can predict time to rebound and precede loss of viral control^22,23^, the association between T cell subsets expressing exhaustion markers and PTC has not been thoroughly evaluated in existing PTC studies. Alternatively, both cytolytic (e.g., directly killing target cells) and non-cytolytic (e.g., transcriptional silencing) CD8+ T cell activity preserved by ART may be essential for PTC^24^. Due to limitations in evaluating HIV-specific CD8+ T cells, it has not been possible to fully define the contributions of CD8+ T cells to PTC in humans. Therefore, a reproducible animal model of PTC would enable experimental interventions like CD8 depletion to be performed, and antigen-specific CD8+ T cells could be evaluated because animals can be MHC-matched.

Rhesus macaques (RMs) infected with simian immunodeficiency virus (SIV) are the most common nonhuman primate model of HIV^25^, yet few become PTCs^26,27^. Definitions of PTC vary between studies but have commonly included maintaining viremia at <10^4^ copies/mL for months to years post ART withdrawal. Initiating RMs on ART within the first five days of infection has been shown to lead to productive infection and result in viral rebound in 100% (9/9) of the RMs within three weeks of ART interruption^13,28^. However, initiating RMs on ART within the first five days of infection can also resemble post-exposure prophylaxis, resulting in RMs that do not rebound after ART interruption but also do not appear to be productively infected^26,29^. Initiating ART within 6-12 days after infection results in productive infection but rarely leads to PTC; Okoye and colleagues found that 32/33 RMs exhibited viral rebound, most within 2-6 weeks after ART withdrawal with plasma viremia >10^4^ copies/mL^26^. Since the SIV RM model of HIV has failed to produce enough PTCs to unravel the underlying mechanisms of PTC, a different model may be required to establish and evaluate PTC.

In the present study, we describe frequent PTC in Mauritian cynomolgus macaques (MCMs). MCMs are an exotic macaque population with limited genetic diversity due to a small number of invasive founder animals colonizing Mauritius within the last several hundred years. The limited genetic diversity extends to the MHC, with only seven MHC haplotypes (M1-M7)^30^ in the entire population. While the M1 and M6 MHC haplotypes are associated with spontaneous SIV control without ART, the M3 MHC haplotype is not^8,30,31^. We initiated ART at two weeks post-infection (wpi) in eight M3+ MCMs that did not have the M1 or M6 haplotypes and discontinued treatment eight months later. For six months after ART withdrawal, plasma viremia in seven of the eight MCMs was either undetectable or only transiently detectable at <10^4^ copies/mL. We find that MCMs had smaller acute reservoirs than similarly infected RMs two wpi. Experimental depletion of CD8α+ cells in all seven PTC MCMs led to viral rebound, demonstrating a role for these cells in maintaining ART-free viral control. M3+ MCMs can now be used to dissect the mechanisms by which small viral reservoirs and CD8α+ cells facilitate PTC.

## Materials and Methods

### Animal care and use

*Animal cohort 1*: Eight male Mauritian cynomolgus macaques (MCMs) purchased from Bioculture Ltd. were housed and cared for by the Wisconsin National Primate Research Center (WNPRC) according to protocols approved by the University of Wisconsin Graduate School Animal Care and Use Committee (IACUC; protocol number G005507). Each animal had at least one copy of the M3 MHC haplotype, and none had the M1 MHC haplotype associated with viral control (Table 1) ^8,30^. All eight MCMs were infected intravenously (i.v.) with 10,000 infectious units (IUs) of barcoded SIVmac239M^32^ and began receiving a daily antiretroviral therapy (ART) regimen consisting of 2.5mg/kg dolutegravir (DTG, ViiV Healthcare, Research Triangle, NC), 5.1mg/kg tenofovir disoproxil fumarate (TDF, Gilead, Foster City, CA), and 40mg/kg emtricitabine (FTC, Gilead) in 15% Kleptose (Roquette America) in water subcutaneously at two wpi. Four of these animals received a therapeutic vaccination regimen during ART described previously^33^. ART was discontinued in all animals after eight months of treatment and the animals were monitored for viral rebound. Six months after ART discontinuation, all eight animals were rechallenged i.v. with 100 TCID50 of non-barcoded SIVmac239. Eight weeks after rechallenge, all eight MCMs received a single 50 mg/kg i.v. infusion of the anti-CD8α mAb MT807R1. The rhesus macaque (RM) IgG1 recombinant Anti-CD8α [MT807R1] monoclonal antibody was engineered and produced by the Nonhuman Primate Reagent Resource (NIH Nonhuman Primate Reagent Resource Cat# PR-0817, RRID:AB_2716320). All animals were necropsied six to eight weeks after CD8α depletion.

**TABLE 1.**
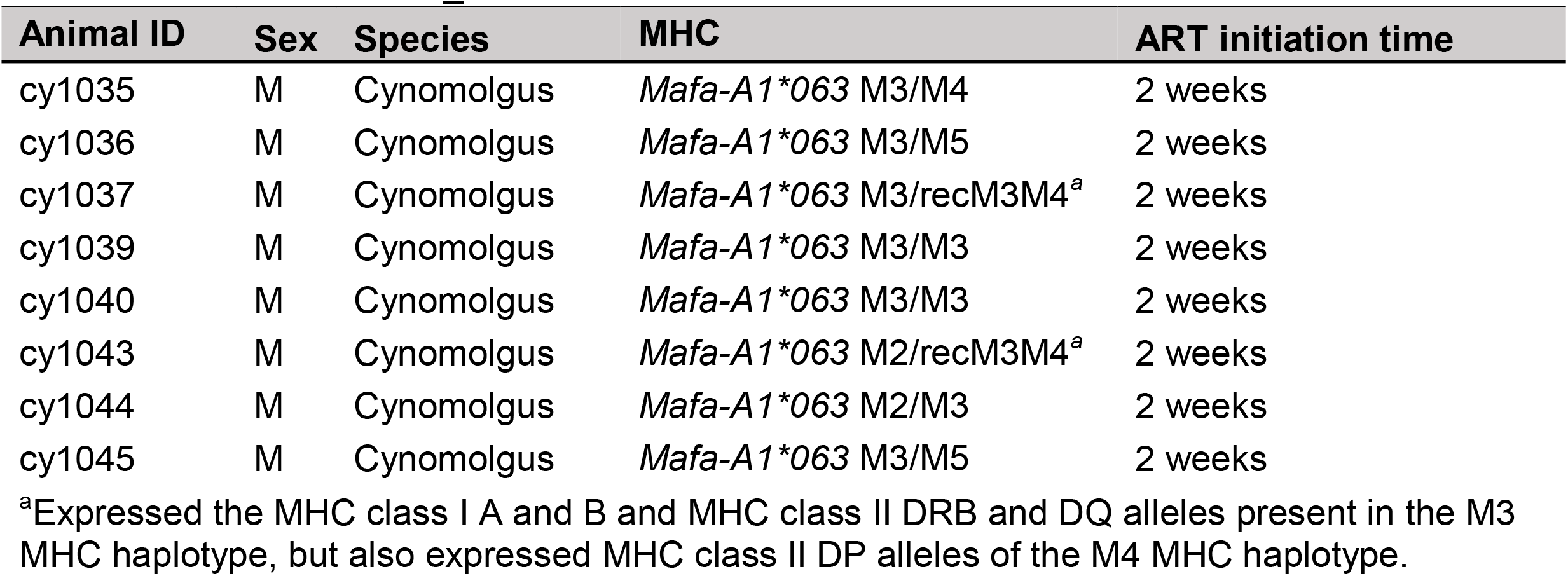
Animals in MCM_2wk cohort

*Animal cohort 2*: Six male MCMs purchased from Bioculture Ltd. were housed and cared for by the WNPRC according to protocols approved by the University of Wisconsin Graduate School (IACUC protocol numbers G005443 and G005507). The MHC haplotypes are summarized in Table 2. These animals had previously been exposed to dengue virus (wild type DENV-1) >5 years before SIV challenge and inoculated i.v. with plasma containing simian pegivirus^34^ ∼5 months after SIV challenge as part of previous projects. All six MCMs were previously inoculated intrarectally with 3,000 TCID50 of SIVmac239. One animal (cy0735) was not productively infected after this inoculation and was therefore rechallenged i.v. with 500 TCID50 of SIVmac239 five weeks after the initial intrarectal challenge. All six MCMs received an ART regimen consisting of 2.5mg/kg DTG (APIChem, Hangzhou, Zhejiang, China), 5.1mg/kg TDF (APIChem), and 50mg/kg FTC (APIChem) in 15% Kleptose (Roquette America) in water subcutaneously daily beginning eight wpi. ART was discontinued in all MCMs after approximately 20 months of treatment and the animals were monitored for viral rebound for 81 days. Animals were transferred to a different study after this time.

**TABLE 2.** Animals in MCM_8wk cohort

| Animal ID | Sex | Species | MHC | ART initiation time |
| --- | --- | --- | --- | --- |
| cy0719 | M | Cynomolgus | *Mafa-A1\*063* M1, M3 | 8 weeks |
| cy0721 | M | Cynomolgus | *Mafa-A1\*063* recM1/M3, M5<sup>a</sup> | 8 weeks |
| cy0723 | M | Cynomolgus | *Mafa-A1\*063* M4, recM3/M1<sup>b</sup> | 8 weeks |
| cy0724 | M | Cynomolgus | *Mafa-A1\*063* M1, M4 | 8 weeks |
| cy0726 | M | Cynomolgus | *Mafa-A1\*063* M2, M3 | 8 weeks |
| cy0735 | M | Cynomolgus | *Mafa-A1\*063* M1, M4 | 8 weeks |

<sup>a</sup>Expressed the MHC class I A allele present in the M1 MHC haplotype, but also expressed the MHC class I B and MHC class II DR, DQ, and DP alleles of the M3 MHC haplotype.
<sup>b</sup>Expressed the MHC class I A and B alleles present in the M3 MHC haplotype, but also expressed MHC class II DR, DQ, and DP alleles of the M1 MHC haplotype.

*Animal cohort 3*: Five Indian RMs (three female and two male) were previously housed and cared for by the WNPRC according to protocols approved by the University of Wisconsin Graduate School (IACUC protocol number G005507). All RMs expressed the *Mamu-A1*001* MHC class I allele and did not express the *Mamu-B*008* or *Mamu-B*017* alleles associated with SIV control^5,6,35^. All RMs were infected i.v. with 10,000 IUs of SIVmac239M as previously described^35^.

### Plasma viral load analysis

Plasma was isolated from undiluted whole blood by Ficoll-based density centrifugation and cryopreserved at -80°C. Plasma viral loads were quantified as previously described^35^. Briefly, viral RNA (vRNA) was isolated from plasma samples using the Maxwell Viral Total Nucleic Acid Purification kit (Promega, Madison WI). Next, vRNA was reverse transcribed using the TaqMan Fast Virus 1-Step qRT-PCR kit (Invitrogen) and quantified on a LightCycler 480 instrument (Roche, Indianapolis, IN). Primers and probes are described in^36^.

### Cell-associated viral RNA (vRNA) analysis

Total RNA was isolated from 1×10^6^ cryopreserved lymph node (LN) mononuclear cells (LNMC) isolated from LN biopsies using the SimplyRNA Tissue kit for the Maxwell RSC instrument (Promega, Madison, WI). RNA was then tested for both SIV *gag* and a cellular gene, β*-actin*, by RT-qPCR. Both assays were performed using the Taqman Fast Virus 1-Step Master Mix RT-qPCR kit (Applied Biosystems Inc., Carlsbad, CA) on an LC96 instrument (Roche, Indianapolis, IN) with identical experimental conditions; 600 nM each primer, 100 nM probe and 150 ng random primers (Promega, Madison, WI). Reactions cycled with the following conditions: 50°C for 5 minutes, 95°C for 20 seconds followed by 50 cycles of 95°C for 15 seconds and 60°C for 1 min. SIV: forward primer: 5’-GTCTGCGTCATPTGGTGCATTC- 3’, reverse primer: 5′-CACTAGKTGTCTCTGCACTATPTGTTTTG-3′, and probe: 5′-6-carboxyfluorescein- CTTCPTCAGTKTGTTTCACTTTCTCTTCTGCG-BHQ1-3’. β-actin: forward primer: 5’- GGCTACAGCTTCACCACCAC-3’, reverse primer: 5’-CATCTCCTGCTCGAAGTCTA-3’, and probe: 5’-6- carboxyfluorescein-GTAGCACAGCTTCTCCTTAATGTCACGC-BHQ1-3’. RNA was quantified for each reaction by interpolation onto a standard curve made up of serial tenfold dilutions of in vitro transcribed RNA.

Because a source of latently infected cells was not readily available, the lower limit of detection was estimated using CD8-macaque PBMCs that were infected with SIVmac239 in vitro at a high multiplicity of infection (MOI). CD8-cells were isolated using a nonhuman primate CD8+ T cell separation kit (Miltenyi Biotec) to collect the unlabeled cell fraction. These cells were confirmed to be SIV+ by qPCR measuring both *gag* and *ccr5* DNA. A dilution series of the SIV+ cells was prepared in 1×10^6^ naïve macaque PBMCs. In these experiments, we can detect as few as 10 infected cells per million PBMCs (unpublished data).

### Quantitative Viral Outgrowth Assays (QVOAs)

QVOAs were modified from previously described work quantifying the HIV latent reservoir^37^. Briefly, peripheral blood mononuclear cells (PBMCs) were isolated from undiluted whole blood by Ficoll-based density centrifugation. Monocytes were removed by adherence via a three-hour incubation at 37ºC. CD4+ T cells were then enriched using a nonhuman primate CD4+ T cell isolation kit (Miltenyi Biotec). 1×10^4^, 1×10^5^, 3×10^5^, 1×10^6^, and/or 2×10^6^ enriched CD4+ T cells were plated in 1-6 replicates per concentration per animal (depending on the available number of cells) at a 1:2 ratio with irradiated (100Gy) CEMx174 cells in R10 (RPMI 1640 medium supplemented with 10% FBS, 1% antibiotic-antimycotic [Thermo Fisher Scientific, Waltham, MA] and 1% L-glutamine [Thermo Fisher Scientific]) in the presence of phorbol myristate acetate (PMA) (0.1μg/mL, Sigma Aldrich, St. Louis, MO) and ionomycin (1μg/mL, Sigma Aldrich) and incubated overnight at 37ºC. The next day, mitogens were removed by washing the contents of each well with R10, and the pellets were resuspended in R15-50 (RPMI 1640 medium supplemented with 15% fetal calf serum, 1% antibiotic/antimycotic 1% L-glutamine, and 50 U/mL of interleukin-2). Fresh CEMx174 target cells were added at a 1:1 ratio with the original plated number of enriched CD4+ T cells and plates were incubated for 2 weeks at 37ºC. After two weeks, supernatant was collected and supernatant viral loads were quantified identically to plasma viral loads, described above. The calculator developed by the Siliciano lab at Johns Hopkins University was used to estimate infectious units per million cells (IUPM) for each animal by combining the number of cells plated, number of replicates at each concentration, and number of positive outcomes (positive supernatant viral loads)^38^.

### Mass Spectrometry Quantification of Plasma ART Concentrations

ART quantification in plasma from SIV+, ART-suppressed MCMs was performed at the Antiviral Pharmacology Laboratory, University of Nebraska Medical Center, applying liquid chromatography tandem mass spectrometry (LCMS) according to previously validated methods^39^. Briefly, DTG concentrations were quantified via a methanolic precipitation extraction with stable-labeled internal standard over the range of 20-10,000 ng/mL. DTG sample extracts were detected via LCMS. TDF/FTC concentrations were quantified via an acidic protein precipitation extraction with stable-labeled internal standards over the range of 10-1,500 ng/mL. TDF/FTC sample extracts were also detected via LCMS.

### DNA extraction

Using the QIAamp DNA Mini Kit (Qiagen), DNA was extracted from previously cryopreserved PBMCs according to the manufacturer’s instructions, except we eluted with 150 μL nuclease-free water. Sample DNA concentrations were measured on a Nanodrop (Thermo Fisher Scientific, MA, USA).

### Intact proviral DNA assay

To quantify frequencies of intact and defective proviral DNA, we used a previously described SIV IPDA^40–42^ with minor modifications^43^. A step-by-step protocol and detailed explanations of our methods are in preparation (Matschke 2023, in prep). To summarize, the SIV IPDA discriminates between intact and defective proviral species by targeting three proviral amplicons across two parallel multiplexed ddPCR reactions, and intact proviruses are defined by non-hypermutated 5’ *pol* and 3’ *env* sequences in the absence of 2-LTR junctions. To normalize proviral frequencies to cell equivalents and correct for DNA shearing, a third parallel ddPCR reaction targets two distinct amplicons within the host ribonuclease P p30 (*rpp30*) gene.

### IFN-_γ_ ELISPOT assays

IFN-γ ELISPOT assays were performed using fresh PBMCs isolated from EDTA-anticoagulated blood by Ficoll-based density centrifugation, as previously described^33^. Peptides (Gag_386-394_GW9, Nef_103-111_RM9, and a Gag peptide pool comprising 15-mer peptides spanning the SIVmac239 Gag proteome, each overlapping by 11 amino acids [NIH HIV Reagent Program, managed by ATCC]) were selected from epitopes restricted by the *Mafa A1*063* MHC class I allele expressed on the M3 MHC haplotype^44^. Assays were performed according to the manufacturer’s protocol, and wells were imaged and quantified with an ELISPOT plate reader (AID Autoimmun Diagnostika GmbH). As described previously^45^, a one-tailed t-test with an α level of 0.05, where the null hypothesis was that the background level would be greater than or equal to the treatment level, was used to determine positive responses. Statistically positive responses were considered valid only if both duplicate wells contained 50 or more spot-forming cells (SFCs) per 10^6^ PBMCs. If statistically positive and ≥50 SFCs per 10^6^ PBMCs, reported values display the average of the two test wells with the average of all four negative control wells subtracted.

### Tetramerization of Gag_386-394_GW9 monomers

Biotinylated monomers were produced by the NIH Tetramer Core Facility at Emory University (Atlanta, GA) using *Mafa*-A1*063 Gag_386-394_GW9 peptides purchased from Genscript (Piscataway, NJ). *Mafa*-A1*063 Gag_386-394_GW9 biotinylated monomers were tetramerized with streptavidin-PE (0.5mg/mL, BD biosciences) at a 4:1 molar ratio of monomer:streptavidin in the presence of a 1x protease inhibitor cocktail solution (Calbiochem, Millipore Sigma). Briefly, 1/5^th^ volumes of streptavidin-PE were added to the monomer for 20 minutes rotating in the dark at 4ºC, and this process was repeated five times.

### Phenotype staining of T cells by flow cytometry

Previously frozen PBMCs isolated from whole blood and previously frozen LNMCs isolated from LN biopsies were used to assess the quantity and phenotype of T cell populations longitudinally. Where tetramer staining was included, cells were thawed, washed with R10, and rested for 30 minutes at room temperature in a buffer consisting of 2% FBS in 1X PBS (2% FACS buffer) with 50nM dasatinib (Thermo Fisher Scientific). Cells were washed with 2% FACS buffer with 50nM dasatinib and incubated with the Gag_386-394_GW9 tetramer for 45 minutes at room temperature. Cells were washed with 2% FACS buffer with 50nM dasatinib and incubated with the remaining surface markers (Table 3) for 20 minutes at room temperature. Where tetramer staining was not included, cells were thawed, washed with R10, and incubated for 20 minutes at room temperature with the surface markers indicated in Table 4. Cells were then washed with 2% FACS buffer and fixed. Where no intracellular staining was performed, cells were fixed for a minimum of 20 minutes with 2% paraformaldehyde and acquired immediately using a FACS Symphony A3 (BD Biosciences). Where intracellular staining was performed, cells were fixed using fixation/permeabilization solution (Cytofix/Cytoperm™ fixation and permeabilization kit, BD Biosciences) for 20 minutes at 4ºC. Cells were washed with cold 1x Perm/Wash™ buffer (Cytofix/Cytoperm™ fixation and permeabilization kit, BD Biosciences) and incubated in 1x Perm/Wash™ buffer containing the CTLA4 antibody (Table 3) for 20 minutes at 4ºC. Cells were washed with 1x Perm/Wash™ buffer and acquired immediately using a FACS Symphony A3 (BD Biosciences). The data were analyzed using FlowJo software for Macintosh (BD Biosciences, version 10.8.0). Cell subpopulations were excluded from analysis when the parent population contained <50 events.

**TABLE 3.**
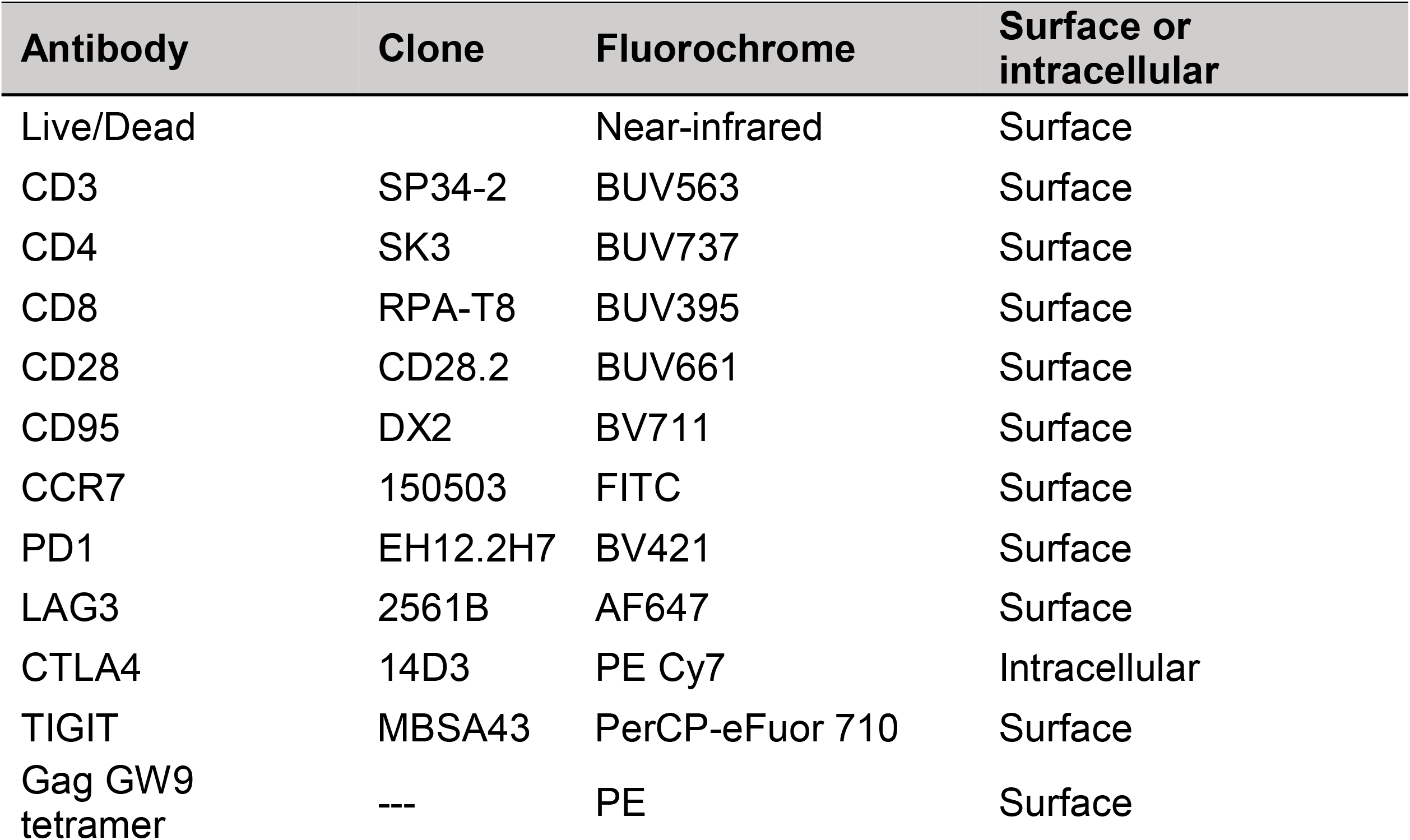
Antibodies used for T cell phenotyping

**TABLE 4.** Antibodies used for T cell and NK cell phenotyping

| **Antibody** | **Clone** | **Fluorochrome** | **Surface or intracellular** |
| --- | --- | --- | --- |
| Live/Dead |  | Near-infrared | Surface |
| CD3 | SP34-2 | AF-700 | Surface |
| CD4 | SK3 | BUV737 | Surface |
| CD8 | DK25 | PE | Surface |
| CD14 | M5E2 | APC-H7 | Surface |
| CD16 | 3G8 | BV786 | Surface |
| CD20 | 2H7 | APC-H7 | Surface |
| CD56 | B159 | FITC | Surface |
| NKG2A/C | Z199 | PE-Cy7 | Surface |

### Intracellular cytokine staining (ICS) assays

ICS assays were performed to characterize T cell polyfunctionality similarly to published work^46,47^. Previously frozen PBMCs isolated from whole blood were thawed, washed with R10, and rested for ∼6 hours in R15 (RPMI 1640 medium supplemented with 15% FBS, 1% antibiotic-antimycotic [Thermo Fisher Scientific], and 1% L-glutamine [Thermo Fisher Scientific]) at 37ºC in 5% CO_2_. After resting, cells were incubated for ∼16 hours at 37ºC in 5% CO_2_ with R15 alone as a negative control, or with the Gag peptide pool described above, at a final concentration of 62.5 μg/mL (0.5 μg/mL of each peptide). Two wells stimulated with 20ng/mL PMA and 1 μg/mL ionomycin were included in each batch of staining as a positive control. 5 μg/mL anti-CD28 and 5 μg/mL anti-CD49d were added during stimulation. After 90-120 minutes, 1 μg/mL Brefeldin A, 2 μM Monensin, and anti-CD107a (Table 5) were added to all cells for the remainder of the stimulation. Following the stimulation, cells were washed with 2% FACS buffer and incubated with the remaining surface markers (Table 5) for 20 minutes at room temperature. Intracellular staining, sample acquisition, and data analysis were performed as described above. All reported values are Gag-specific responses with media controls background subtracted. When Gag-specific responses were not higher than the background, values are reported as zero.

**TABLE 5.** Antibodies used for ICS assay

| **Antibody** | **Clone** | **Fluorochrome** | **Surface or intracellular** |
| --- | --- | --- | --- |
| Live/Dead |  | Near-infrared | Surface |
| CD3 | SP34-2 | BUV563 | Surface |
| CD4 | SK3 | BUV737 | Surface |
| CD8 | RPA-T8 | BUV395 | Surface |
| CD107a | H4A3 | PE | Surface |
| IFN-γ | 4S.B3 | FITC | Intracellular |
| TNF-α | MAB11 | AF700 | Intracellular |
| IL-2 | MQ1-17H12 | BV605 | Intracellular |
| MIP1β | D21-1351 (RUO) | BV421 | Intracellular |

### Enzyme-Linked Immunosorbent Assays (ELISAs)

Uncoated 2HB 96-well plates (Thermo Fisher Scientific) were coated with 0.5 μg/mL SIVmac239 gp120 (Immune Technology, New York, NY) in coating buffer (2.25mg/mL Na_2_CO_3_ and 4.395mg/mL NaHCO_3_ in distilled H_2_O). Plates were incubated overnight at 4ºC. The next day, plates were washed with wash buffer (1X PBS with 0.05% V/V TWEEN-20 [Thermo Fisher Scientific]) and blocked in a buffer containing 1X PBS supplemented with 10% FBS for one hour at room temperature. After blocking, plates were washed and serial dilutions of plasma were added for 90 minutes at room temperature. Plasma samples were heat-inactivated at 56ºC for 30 minutes, prior to use. Serial dilutions of anti-SIV gp120 monoclonal antibody (B404, NIH HIV Reagent Program, Division of AIDS, NIAID, NIH) were plated as a positive control. Next, plates were washed with wash buffer and anti-monkey IgG HRP (SouthernBiotech, Birmingham AL) was diluted 1:10,000 in blocking buffer and added to each well. Plates were incubated for one hour at room temperature. Plates were then washed with wash buffer and TMB substrate (3,3’5,5’ – tetramethylbenzidine, Thermo Fisher Scientific) was added to each well. After a 15 minute incubation, HCl (1N) was added to each well to stop the reaction. ELISA plates were immediately read using a GloMax®-Multi Detection System microplate reader (Promega, Madison WI) at 450 nm absorbance.

### RNA sequencing (RNAseq)

Previously cryopreserved PBMCs were thawed and CD8+ T cells were enriched by negative selection using a nonhuman primate CD8+ T cell isolation kit (Miltenyi Biotec). RNA was isolated from the enriched CD8+ T cells by TRIzol/phenol chloroform extraction with DNase treatment. RNAseq was performed by the University of Wisconsin Biotechnology Center Gene Expression Center. Briefly, the library was prepared by synthesizing cDNA via reverse transcription and amplifying cDNA by long-distance PCR using the Takara SMARTer v4 kit (Takara Bio USA, San Jose, CA). 2×150bp sequencing was performed on an Illumina NovaSeq6000 instrument. Mean +/- SD sequencing depth was 265.75M +/- 45.59M total reads. RNA sequencing data were deposited in the NCBI’s Gene Expression Omnibus and are publicly available under GEO accession GSE225770.

### RNAseq analyses

BCL to FASTQ conversion was performed, then FASTQ files were used as input into the NextFlow (v22.04.5^48^) pipeline nf-core/rnaseq pipeline (v3.8.1^49,50^) using *Macaca fascicularis* genome version 6.0.107 (MacFas6) for alignment and annotation. Mean +/- SD alignment rate was 82.4% +/- 1.0%. Subsequent analysis was performed on the Salmon gene count matrix, with removal of genes where the total count across samples was less than 10, as well as redundant gene_name rows (Y_RNA) and all poorly annotated ENSMFAG# format genes. EBSeq^51,52^ was used to determine statistical significance, due to better model fit than DESeq2, and was run for 2 condition comparison on median normalized counts data for a total of 5 iterations. Genes with a false discovery rate < 0.05 were considered significant.

### Statistical analyses

For statistical analyses in which animal groups were being compared to each other at the same time point, Mann-Whitney U tests were performed. Wilcoxon signed-rank tests were used to compare the same individuals at two time points. Unpaired t-tests with Welch’s correction were performed to compare animal groups of different species. Paired t-tests were performed to compare two time points from the same animal group. Chi-square tests were performed to compare frequency distributions of vDNA types in IPDAs. When three animal groups were compared across two time points, two-way ANOVA tests were performed with Šídák correction for multiple comparisons. Correlation analyses were performed via nonparametric Spearman correlation. R^2^ values and best-fit trend lines were computed using simple linear regression. All statistical analyses except those for RNAseq were calculated in GraphPad Prism.

## Results

### Prolonged control of SIV replication after ART withdrawal in M3+ MCMs initiated on ART at two wpi

We previously infected eight MCMs intravenously (i.v.) with 10,000 infectious units (IUs) of SIVmac239M and started them on antiretroviral therapy (ART) two weeks later (Fig. 1a; MCM_2wk cohort). None of these MCMs possessed the M1 MHC haplotype associated with SIV control (Table 1)^8,30^. We included animals possessing at least one copy of the M3 MHC haplotype that is not associated with spontaneous SIV control. Four animals (denoted by open symbols) received therapeutic vaccinations described elsewhere^33^, and four animals received only ART. All eight animals received ART for approximately eight months before ART was interrupted. Up to 23 weeks after ART withdrawal, only one animal (cy1043) displayed sustained viral rebound beginning nine weeks after ART withdrawal (Fig. 1b). The other seven animals maintained undetectable or transiently detectable viremia at <10^4^ copies/mL.

**Fig. 1.**
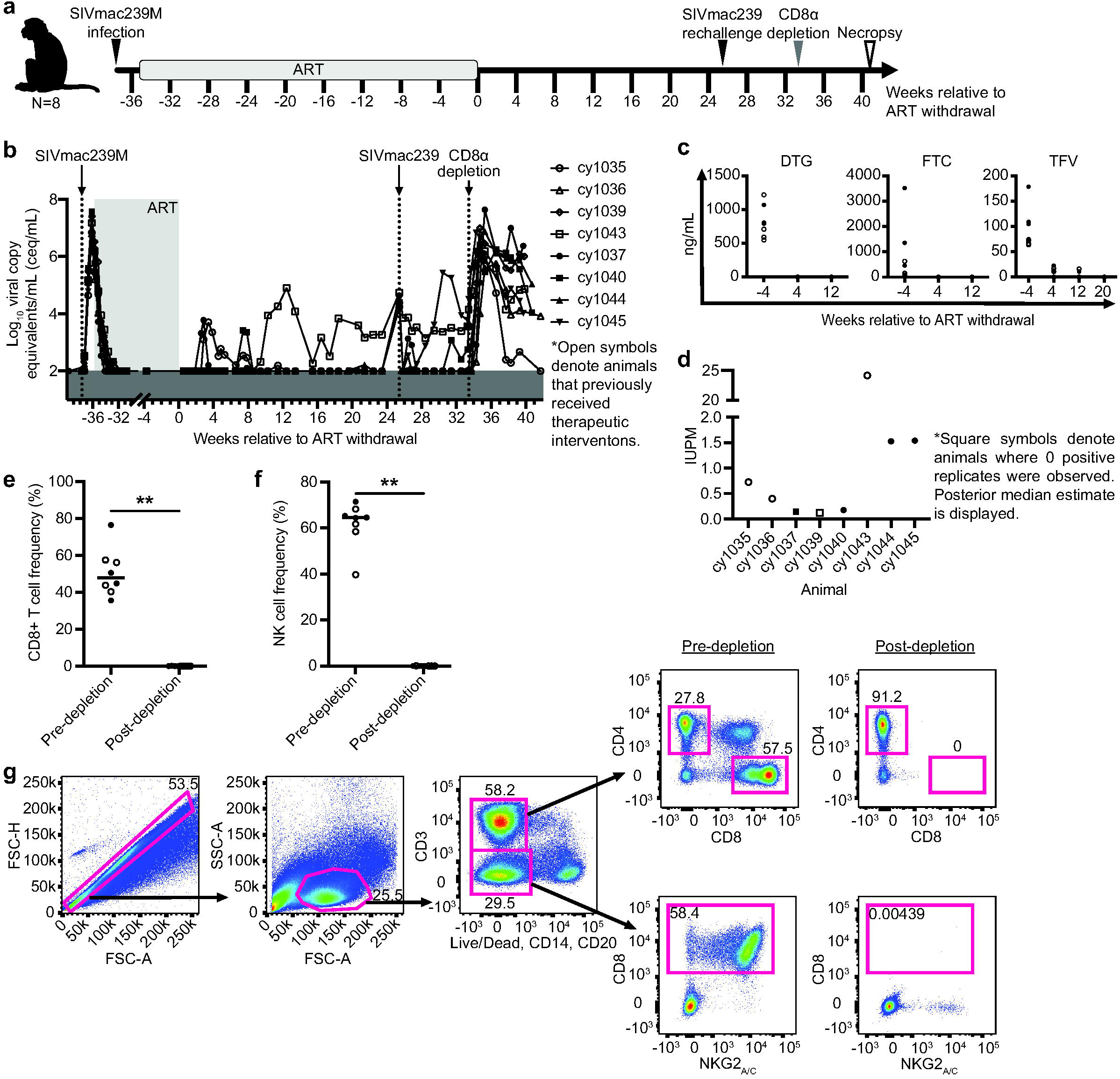
Experimental design, ART drug concentrations, and immunological and virological analyses. **a** Study design depicting the timeline of SIVmac239M infection, ART initiation, ART withdrawal, SIVmac239 rechallenge, CD8α depletion, and necropsy. Animals that previously received therapeutic interventions are displayed as open symbols. **b** Individual plasma viral loads from SIVmac239M infection through ART initiation, ART withdrawal, SIVmac239 rechallenge, CD8α depletion, and necropsy. Viral loads are displayed as log_10_ceq/mL. **c** Plasma concentrations of DTG (left), FTC (center), and TFV (right) during ART treatment (the -4 time point) and after ART withdrawal. Amounts BLQ are displayed as zero ng/mL. **d** IUPM for each animal ∼19-21 weeks after ART withdrawal determined by QVOA. Square symbols denote animals where zero positive replicates were observed; posterior median estimate is displayed. **e**-**f** The frequency of CD8+ T cells (**e**) and NK cells (**f**) before and 10 days after CD8α depletion. Results are displayed for each animal individually with the lines at the median. ** *P*=0.0078. P values were calculated using a Wilcoxon signed-rank test. **g** Representative gating strategy used to evaluate CD8+ T cells (top images) and NK cells (bottom images) before (left) and 10 days after CD8α depletion (right).

To investigate whether PTC was due to the residual presence of ART drugs, we measured plasma concentrations of DTG, TFV, and FTC. All three drugs were detectable in the normal range seen in humans^53,54^ during ART and dropped below the limit of quantification (BLQ) by one to five months post-ART withdrawal (Fig. 1c).

Next, we performed quantitative viral outgrowth assays (QVOAs) with PBMCs collected ∼19-21 weeks after ART withdrawal to determine if replication-competent SIV remained. SIV *gag* was detected by qRT-PCR in at least one replicate in six of the eight animals (Fig. 1d), indicating that reactivatable SIV was present.

### Rechallenge with an isogenic virus is largely contained in PTCs

Approximately 25 weeks after ART withdrawal, all eight MCMs were rechallenged i.v. with SIVmac239 to determine whether they possessed immune responses that could protect from rechallenge. SIV *gag* RNA was detected in plasma collected within 70 minutes of administering SIVmac239, demonstrating that SIV was infused. Four animals (cy1037, cy1040, cy1043, and cy1045) either rebounded or had transient detectable blips in viremia within six weeks of rechallenge while viremia in the other four animals (cy1035, cy1036, cy1039, and cy1044) remained undetectable (Fig. 1b). Sequencing plasma virus during this period revealed that circulating virus on the day of challenge was almost exclusively derived from SIVmac239, while >90% of the virus detectable in the blips contained a molecular barcode derived from SIVmac239M (data not shown, Moriarty in prep). IFN-γ ELISPOT assays indicate that after rechallenge, the animals with undetectable viremia had more positive T cell responses (measured by the number of spot-forming cells [SFCs] per 1×10^6^ PBMCs) to a pool of Gag peptides, Gag_386-394_GW9, and Nef_103-111_RM9 than animals with detectable post-rechallenge viremia (Supplemental Fig. 2). These results suggest that post-rechallenge viral control may be associated with higher SIV-specific cellular immune responses.

### CD8_α_+ immune cells mediate PTC

Eight weeks after SIVmac239 rechallenge, we transiently depleted CD8α+ cells *in vivo* from all eight animals of (Fig. 1a) by intravenously administering the MT807R1 CD8α-depleting antibody. We observed a ∼3-6 log_10_ increase in SIV viremia within ten days of administration of the CD8α-depleting antibody (Fig. 1b), corresponding with a significant decline in the frequency of CD8+ T cells and NK cells (Fig. 1e-g). These results suggest that post-ART immune control was mediated by CD8α+ cells.

### MCMs have smaller total and intact viral reservoirs than RMs at two wpi

It is widely believed that smaller viral reservoirs increase the likelihood of PTC ^55–58^. Much of the integrated vDNA in HIV+ patients is defective^59^ but assays that can quantify the replication-competent reservoir may better predict the likelihood of viral rebound after ART withdrawal^42^. Intact proviral DNA assays (IPDAs) estimate the number of cells containing vDNA and distinguish defective proviruses from intact proviruses, which are likely replication-competent^42^. We used IPDAs to compare the number of intact and total vDNA copies per 1×10^6^ cell equivalents between MCMs at the time of ART initiation, and a separate cohort of RMs infected with SIVmac239M by the same dose and route^35^. RMs initiated on ART ∼one to four weeks post SIV exhibit prompt viral rebound after ART interruption^18,26,47,55,60^, but it is unknown if differences in reservoir sizes between RMs and MCMs account for the different rebound kinetics after ART withdrawal.

We found that RMs had significantly more total and intact vDNA than MCMs at two weeks post-infection (wpi; Figs. 2a and b, respectively). The distribution of vDNA species within the total vDNA compartment was similar between RMs and MCMs (Fig. 2c). Even though the RM viral loads at two wpi were approximately 1 log_10_ higher than MCMs (Fig. 2e), both species had similar peak viral loads (Fig. 2d). We concluded that overall acute SIV plasma viremia was similar in both MCMs and RMs, but there are approximately 10-fold more cells containing intact proviral DNA in RMs compared to MCMs. Unfortunately, the RMs never received ART and could not be compared to the MCM cohort after this time point.

**Fig. 2.**
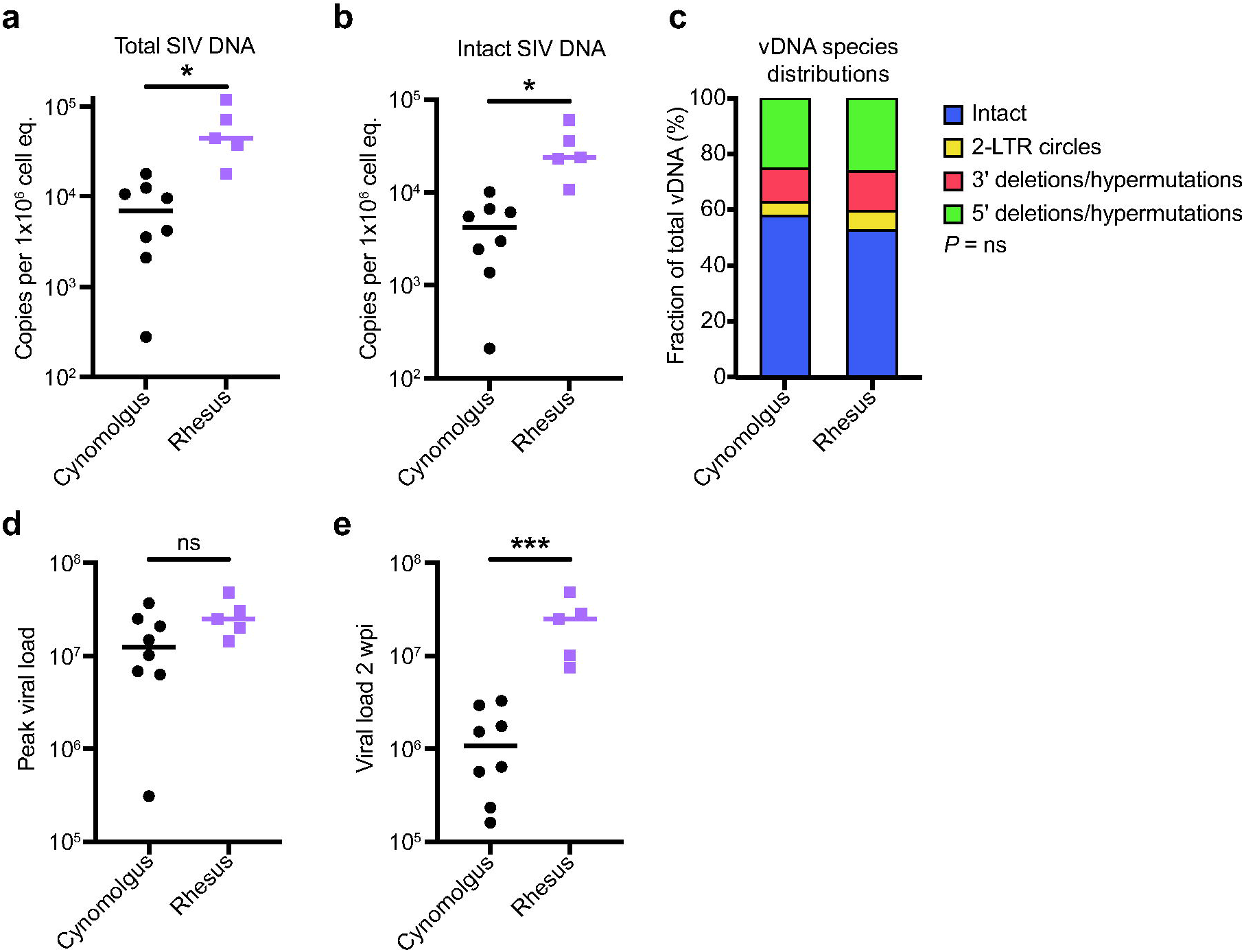
Total and intact proviral DNA in the peripheral blood is lower in cynomolgus macaques than in rhesus macaques during acute SIV infection. **a-b** Total (**a**) and intact (**b**) vDNA per million cell equivalents in PBMCs from the eight cynomolgus macaques (black) or five rhesus macaques (purple) collected two wpi. **c** The distribution of vDNA (vDNA) types. **d-e** Peak viral load (**d**) and viral load 2wpi (**e**) for the rhesus and cynomolgus macaques. ns P≥0.05, * P≤0.05, ** P≤0.01, *** P≤0.001. Unpaired t-tests with Welch’s correction were performed for **a, b, d**, and **e**, and a Chi-square test was performed in **c**. Median and individual values are shown in **a, b, d**, and **e**, and mean fractions are shown in **c**

### A longer duration of SIV infection prior to ART does not increase the frequency of cells with intact proviral DNA but increases the likelihood of viral rebound

We wanted to determine if MCMs were predisposed to having fewer cells with intact proviral DNA or if the frequency of these cells increased with a longer duration of infection prior to ART initiation. We performed IPDAs on samples from a second cohort of six MCMs that were intrarectally infected with 3,000 TCID50 of SIVmac239 and started on ART at eight wpi (Fig. 3a; MCM_8wk cohort). Unlike the MCM_2wk cohort, a subset of these animals expressed the M1 MHC haplotype associated with SIV control (Table 2)^8,30^. We were surprised that the number of intact proviral DNA copies per million cell equivalents was slightly lower in the MCM_8wk cohort at 8wpi (Median=3.1×10^3^ copies per 1×10^6^ cell equivalents) compared to the MCM_2wk cohort at 2wpi (Median=4.3×10^3^ copies per 1×10^6^ cell equivalents). Unfortunately, PBMCs collected at 2wpi from the MCM_8wk cohort were unavailable, so we could not make a longitudinal comparison within the animal group.

**Fig. 3.**
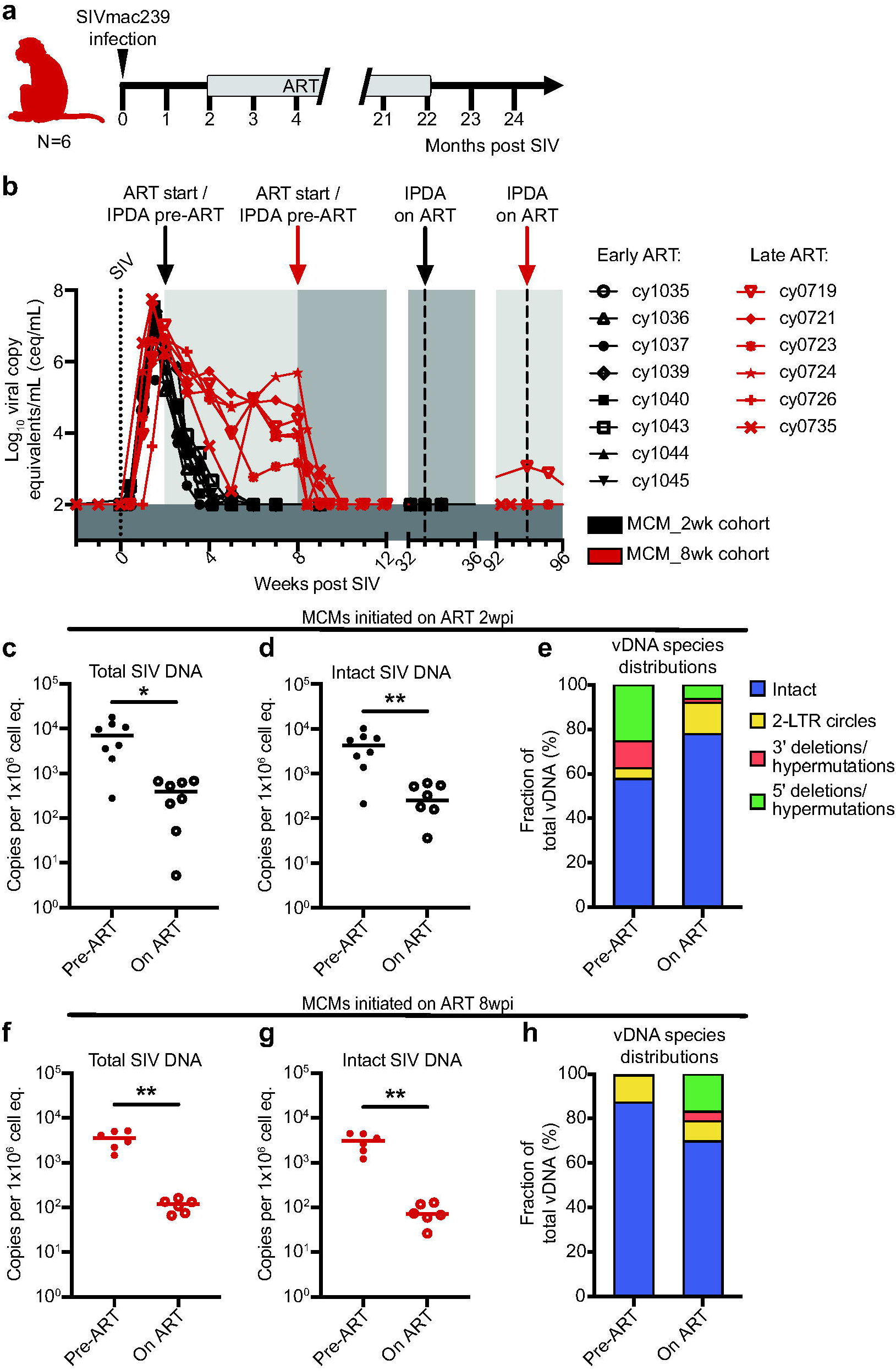
Experimental design for the MCM_8wk cohort and total and intact proviral DNA in the peripheral blood of the MCM_8wk and MCM_2wk cohorts. **a** Study design depicting the timeline of SIVmac239 infection, ART initiation, and ART withdrawal for the six MCM_8wk animals. **b** Individual plasma viral loads depicting the timing of ART initiation and time points for IPDA vDNA analyses for all 14 MCMs. **c**-**e** Total vDNA (**c**), intact vDNA (**d**), and the distribution of vDNA types (**e**) in the eight MCM_2wk animals before (2wpi, pre-ART) and 31 weeks after ART initiation (33wpi, on ART). **f**-**h** Total vDNA (**f**), intact vDNA (**g**), and the distribution of vDNA types (**h**) in the six MCM_8wk animals before (8wpi, pre-ART) and 86 weeks after ART initiation (94wpi, on ART). * P≤0.05, ** P≤0.01. Paired t-tests were performed for **c, d, f**, and **g**. Median and individual values are shown in **c, d, f**, and **g**, and mean fractions are shown in **e** and **h**

The size of the viral reservoir decays during ART, even though it is not eliminated^14,61^. We wanted to determine if the number of intact proviral DNA copies per million cell equivalents declined in MCMs during ART. We performed IPDAs on PBMCs collected during chronic ART, but prior to ART withdrawal (Fig. 3b). In both MCM cohorts, there was a significant decline in total and intact vDNA (Fig. 3c, d, f, g). Statistical analyses were not performed on the vDNA species distributions because the on ART distributions represent a small number of sequences near the limit of detection (Fig. e, h).

### Despite similar reservoir size, viral rebound can occur in MCMs initiated on ART at eight wpi

We hypothesized that the MCM_8wk cohort would also exhibit PTC because they had fewer cells containing intact proviral DNA than the MCM_2wk cohort. Thus, we discontinued ART for these six animals after ∼20 months of ART and viral loads were monitored for 12 weeks. Four of the six MCM_8wk cohort animals exhibited sustained post-ART viremia (>5×10^2^ copies/mL) within four weeks of ART withdrawal and the other two animals exhibited transiently detectable blips in viremia 6-12 weeks after ART withdrawal (Fig. 4a). This contrasted the MCM_2wk cohort, where only one of eight animals developed sustained plasma viremia beginning nine weeks post ART withdrawal, three exhibited transient viral blips, and the remaining four maintained undetectable viremia (Fig. 4b).

**Fig. 4.**
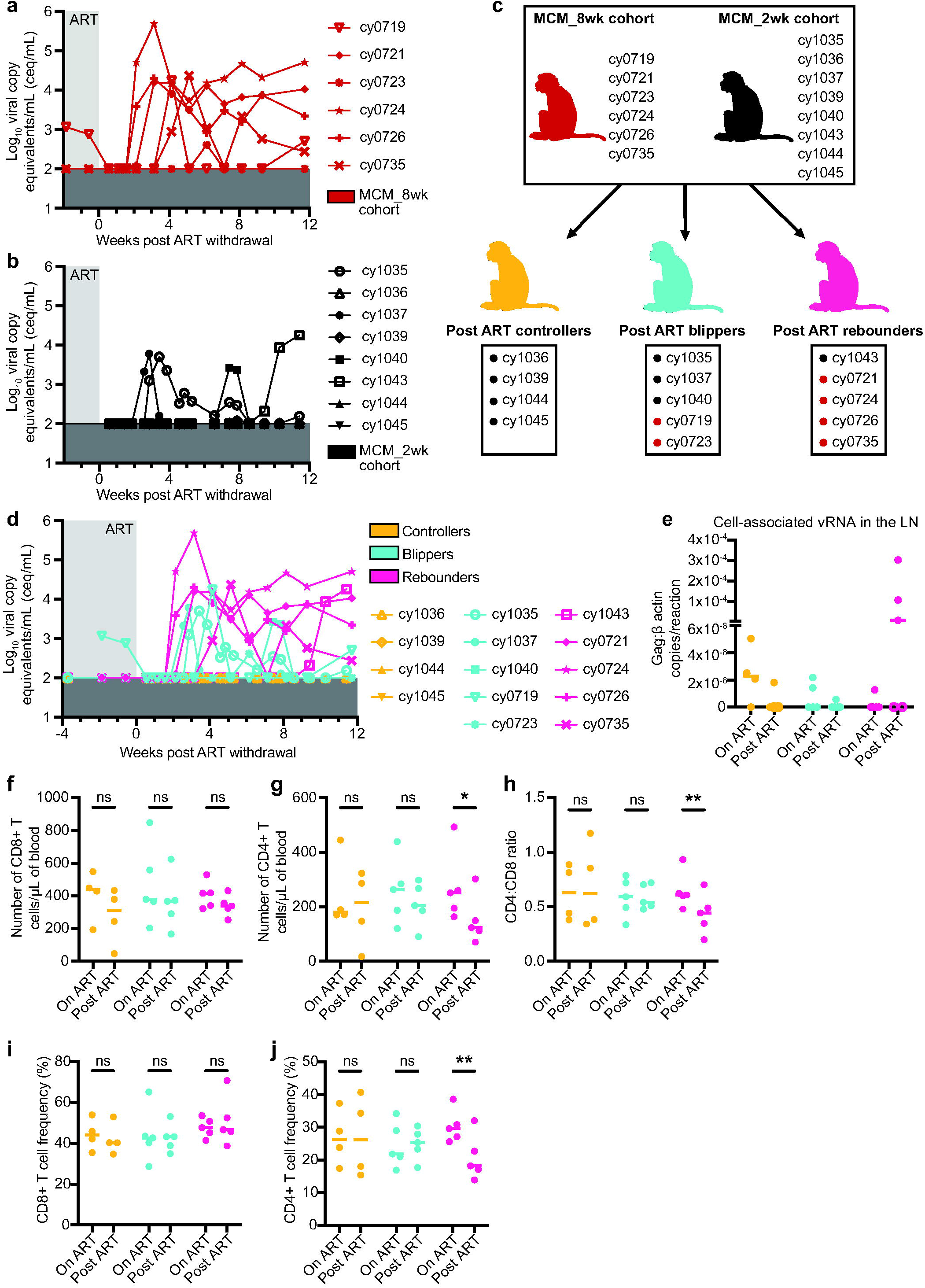
Rearrangement of both animal cohorts into post-ART SIV controllers, blippers, and rebounders, and differences in CD4+ and CD8+ T cell subsets. **a-b** Individual plasma viral loads for the MCM_8wk cohort (**a**) and the MCM_2wk cohort (**b**) from two weeks before ART withdrawal to 12 weeks post ART withdrawal. Viral loads are displayed as log_10_ceq/mL. **c** Regrouping of animal cohorts into animals that, after ART withdrawal, controlled (orange), blipped (teal), or fully rebounded (pink). **d** Combined post-ART viral loads for the controllers, blippers, and rebounders. **e** Cell-associated vRNA in the LNs of the animals during ART treatment (on ART) and 3 weeks post ART withdrawal (post ART). Results are displayed for each animal as the ratio of Gag:β actin per reaction with the lines at the median. Open circles denote samples where Gag was detectable but not quantifiable. **f** Number of CD8+ T cells per µL of blood, **g** number of CD4+ T cells per µL of blood, **h** CD4:CD8 ratio, **i** CD4+ T cell frequencies, and j CD8+ T cell frequencies in the controllers (black), blippers (teal), and rebounders (pink) during ART treatment (on ART) and 12 weeks post ART withdrawal (post ART). Results are displayed for each animal individually with the lines at the median. * P≤0.05, ** P≤0.01. P values were calculated using two-way ANOVA tests with Šídák correction for multiple comparisons.

We re-categorized all 14 MCMs during the 12 weeks after ART interruption (eight from the MCM_2wk cohort and six from the MCM_8wk cohort) as post-ART controllers (maintained undetectable viremia), blippers (had transient viremia above the limit of detection), or rebounders (exhibited sustained viremia above 5×10^2^ copies/mL without re-control) (Fig. 4c). Combined plasma viral loads are displayed in Fig. 4d. Across all animals, there were no significant differences in the cell-associated viral RNA (vRNA) in total lymph node (LN) homogenates either during ART or three weeks after ART withdrawal (Fig. 4e). There were also no differences in the plasma anti-SIVmac239 gp120 IgG antibodies between the groups during ART, 10 days, or 80 days after ART withdrawal (Supplemental Fig. 3).

### PTC in MCMs is correlated with preserved CD4+ frequencies and lower frequencies of CD8+ T cells expressing exhaustion markers

We hypothesized that high CD8+ T cell frequencies may have contributed to PTC, but we found no differences in CD8+ T cell numbers or frequencies in PBMCs from the controllers, blippers, or rebounders during ART or 12 weeks post ART withdrawal (Fig. 4f, i). CD4+ T cell numbers, frequencies, and the CD4:CD8 ratio significantly decreased in the rebounders after ART withdrawal (Fig. 4g, h, j). This was not surprising because increased viral replication is associated with CD4+ T cell decline in viral rebounders^62^. CD4+ T cell frequencies before ART withdrawal were similar between groups and did not predict PTC. To test the hypothesis that the PTCs had T cells with greater functional capabilities than the blippers or rebounders, we measured the frequency of CD4+ and CD8+ T cells expressing CD107a, TNF-α, and IFN-γ after Gag peptide pool stimulation in vitro (Supplemental Fig. 4). Surprisingly, we found no differences in the frequencies of CD4+ or CD8+ T cells expressing combinations of one, two, or three cytokines/cytolytic markers during ART or 12 weeks post ART withdrawal (Supplemental Fig. 4c, d). We also measured differential gene expression in CD8+ T cells from four PTCs and four rebounders during ART (time point 1) and after ART withdrawal (time point 2; 12 weeks post ART withdrawal in the controllers or the last time point with undetectable viremia prior to rebound in the rebounders). While there were marked changes in gene expression between time points, these changes were similar among controllers and rebounders. (Supplemental Fig. 5).

Because T cell exhaustion has been implicated in HIV/SIV disease progression and loss of viral control (reviewed in^63^), we evaluated T cell subsets expressing exhaustion markers (Supplemental Figs. 6-8). The frequency of bulk CD8+ T cells, T_CM_, T_TM_, and T_EM_ expressing CTLA4 significantly declined from on ART to 12 weeks post ART withdrawal in the blippers and controllers, and significantly positively correlated with viral load after ART withdrawal across all groups (Supplemental Fig. 6). The frequency of Gag-specific (Gag_386-394_GW9 tetramer+) CD8+ T cells expressing LAG3 and CTLA4 also significantly declined in the controllers after ART withdrawal (Supplemental Fig. 7b-c). Lastly, we observed a reduced frequency of CD4+ T_CM_ and T_EM_, and CD8+ T_CM_ in the controllers expressing PD1 after ART withdrawal (Supplemental Fig. 8).

## Discussion

Here, we provide the first example of consistent PTC of SIV in a cohort of M3+ MCMs infected with SIVmac239M and initiated on ART at two wpi. Although ART drugs had been cleared systemically (Fig. 1c) and replication-competent virus was present (Fig. 1d), seven of the eight M3+ MCMs had undetectable plasma viremia over six months after ART withdrawal. Our results implicate both CD8α+ cells and the formation of unusually small viral reservoirs in post-ART SIV suppression.

Multiple lines of evidence, including post-CD8 depletion viral rebound and similar CD8+ T cell frequencies in PTCs and rebounders (Figs. 1, 4), suggest that other differences in CD8+ T cells in the MCMs contributed to differing virological outcomes. Elevated expression of immune checkpoint inhibitors (ICIs) like PD1, CTLA4, and LAG3 on T cells coincides with progressive loss of effector function, and this dysfunction has been implicated in the loss of viral control and disease progression (reviewed in^63^). Because the frequency of bulk and SIV-specific T cell subsets expressing ICIs declined after ART withdrawal in PTCs but not in animals that rebounded, reduced immune exhaustion may contribute to PTC. CD8+ T cells may also suppress virus replication through non-cytolytic mechanisms, such as suppressing virus transcription^24^. Our observation that three MCMs maintained undetectable plasma viremia between ART interruption and CD8 depletion indicates that either the CD8+ T cells were effectively killing infected CD4+ T cells without evidence of virus replication or SIV transcription was blunted.

PTC may be associated with events of acute infection that are not restored by ART-mediated virus suppression^64^. The earlier ART is initiated during acute infection, the higher the likelihood of later immunologic control after ART withdrawal^2,4^. This is partially attributed to limited reservoir establishment. Individuals with smaller viral reservoirs are overrepresented among identified PTCs, but the contribution of reservoir size to predicting PTC and time to rebound is unclear^2,58,65^. We found the total and intact vDNA reservoirs were approximately 10-fold lower in MCMs than RMs two wpi (Fig. 2), suggesting that MCMs naturally develop smaller viral reservoirs than RMs. Within MCMs, we could not distinguish PTCs from rebounders by the size of the intact or total vDNA reservoirs on ART or after ART withdrawal (Figs. 3, 4). The rebounders may have had more vDNA in tissue reservoir sites like the gut that comprise the majority of the viral reservoir, but these sites were not sampled. This possibility should be examined in future MCM studies. Collectively, MCMs consistently develop small viral reservoirs, but this is not sufficient to predict PTC.

PTC in humans is rare (typically estimated at 4-16% of HIV+ individuals^1,2,4^), and only 22% of the PTCs identified in the CHAMP study (1% of all individuals examined) maintain PTC for ≥5 years^1^. A relevant animal model would enable the immunological and virological correlates of PTC to be characterized. Consistent PTC in MCMs without known protective MHC alleles in this study aligns with the observation that HIV+ PTCs are not enriched for protective MHC alleles^2,8,9,30^. PTC in MCMs could thus be a good model for PTC in humans. Recently, Sharaf and colleagues were the first to use IPDA assays to measure the viral reservoir size in PTCs and reported that PTCs had nearly ten-fold less intact proviral DNA genomes than non-controllers^65^. MCMs had approximately ten-fold less total and intact vDNA than RMs. Bender and colleagues found that HIV+ humans had ten-fold less intact vDNA than RMs in the peripheral blood^40^. The viral reservoir size of SIV+ MCMs is, therefore, more similar to HIV+ humans than SIV+ RMs. SIV+ MCMs recapitulate multiple aspects of PTC observed in humans, potentially making them a better model of PTC than RMs.

Most RM ART interruption studies identify zero PTCs^18,26,47,55,60^. While the definition of PTC varies dramatically between studies^10^, many definitions of PTC (e.g., maintain plasma viremia <10^4^ copies/mL^27^) are lenient enough to include at least nine of the 14 MCMs described here. Our results indicate that PTC is much more common in M3+ MCMs than RMs. Even the four MCM_8wk animals that rebounded within four weeks of ART interruption had peak rebounf viremia of only 4.2-5.7 log_10_ceq/mL (Fig. 4a). In contrast, RMs initiated on ART in similar timeframes consistently rebound within two weeks of ART interruption with peak rebound viremia of ∼4-7 log_10_ceq/mL^18,26,47,55^. Thus, even when MCMs exhibit viral rebound, viremia is delayed and of a smaller magnitude compared to RMs. Although MCMs can develop high peak and set point viremia akin to that of RMs during untreated SIV infection^30^, MCMs and RMs have distinct post-ART rebound kinetics. We propose that other differences in viral load kinetics and pathogenesis between these species may, therefore, also exist.

It would be interesting to directly compare MCMs to RMs in an ART interruption study. While acute CD8+ T cells do not impact reservoir establishment in RMs^18^, it is possible that CD8+ T cells impair reservoir establishment in MCMs. Depletion of CD8+ T cells during acute SIV infection in MCMs could help determine if CD8+ T cells are required for reservoir formation, virus suppression under ART, and later PTC. Even though similar depletion studies were performed in RMs^18^, acute depletion studies in this novel MCM model of PTC may identify key features of immune-mediated virus suppression that were otherwise confounded by the larger viral reservoir and rapid viral rebound in RMs. CD4+ T cells in RMs may also be more susceptible to SIV infection than CD4+ T cells in MCMs, which would affect reservoir formation and size. Therefore, future studies directly comparing RMs and MCMs would be important to better understand early reservoir formation, PTC, and which macaque models are best for studying a functional HIV cure.

The limited genetic diversity of MCMs permits a more precise evaluation of the genetic basis of CD8+ T cell responses associated with virus suppression than is possible in the more genetically diverse RMs. For example, MCMs, like humans, express a lower number of MHC-E alleles (three and two, respectively) than RMs (25), but MHC-E is functionally conserved among MCMs, humans, and RMs^66,67^. CD8+ T cells recognize pathogen-derived peptides in the context of MHC-E and can protect against SIV^68^. Antigen-specific MHC-E-restricted CD8+ T cells can perform antiviral functions like secreting antiviral cytokines and killing infected cells^69^, so these responses may contribute to PTC in MCMs but not RMs. This should be evaluated in ART interruption studies of MCMs with different genetic backgrounds to identify the contributions of protective and non-protective MHC haplotypes to CD8+ T cell responses in the context of PTC.

It would also be beneficial to attempt to “break” PTC in MCMs to characterize the contribution of CD8+ T cell function to PTC. Initiating ART 2wpi in MCMs may be the “goldilocks” time to allow antiviral immunity to develop but not become dysregulated. Therefore, if ART was initiated later in chronic infection (i.e., six months post SIV), then viral evolution, reservoir expansion, and immune exhaustion may prevent PTC. Alternatively, MCMs could be infected with an SIV strain containing mutations in CD8+ T cell epitopes^70^ to determine if acute SIV-specific CD8+ T cells are required for subsequent PTC. An epitope knock-out SIV strain could exploit the limited genetic diversity of MCMs to understand if acute antigen-specific CD8+ T cells contribute to PTC. It is unclear whether differences in acute bulk or SIV-specific CD8+ T cells contributed to the observed PTC. Preventing PTC in MCMs may help explain why MCMs become PTCs.

One drawback of this study was that some of the MCM_2wk animals received therapeutic interventions described elsewhere^33^, but these interventions did not impact viral rebound. Further, the MCM_8wk cohort (Fig. 3) was an imperfect match to the MCM_2wk cohort (Fig. 1) due to differences in MHC genetics, route of SIV challenge, timing of ART initiation, and duration of ART suppression. However, all MCMs had similar amounts of total and intact vDNA in the PBMCs before ART initiation and similar amounts of cell-associated vRNA in the LNs, indicating that these animals were an appropriate comparator group (Figs. 3d, 4e). Despite these limitations, it is clear that MCMs can become PTCs when initiated on ART at two wpi. We can now use this model to further evaluate the predictors of PTC in additional animal studies.

In sum, we have identified a novel model of PTC using M3+ MHC haplomatched MCMs. Of the 14 animals in this study, four maintained undetectable viremia, five had transient blips in viremia, and five exhibited sustained viremia after ART withdrawal. Interruption of ART in SIV+ MCMs leads to very different rebound kinetics (i.e. lower post-ART peak and set point viral load and longer time to rebound) compared to RMs described in the literature with similar infection histories^26^. Future post-ART SIV remission studies using this novel MCM model of PTC may inform the development of therapeutic interventions.

## Author contributions

OEH, ALE, DHO, and SLO contributed to the conception and design of the experiments. CVF, JDL, TCF, MRR, and SLO provided supervision and reviewed data. OEH, LMM, AJB, AJW, AMW, and LCW conducted experiments. OEH, LMM, and JDL analyzed the data. JDL and RVM contributed to data availability. BFK and DHO provided key resources. OEH and SLO wrote the manuscript.

## Acknowledgments

Antiviral drugs were generously provided by Gilead (TDF and FTC) and ViiV Healthcare (DTG). The following reagent was obtained through the NIH HIV Reagent Program, Division of AIDS, NIAID, NIH: Peptide Pool, Simian Immunodeficiency Virus (SIV)mac239 Gag Protein, ARP-12364, contributed by DAIDS/NIAID. The Anti-CD8α [MT807R1] antibody used in this study was provided by the NIH Nonhuman Primate Reagent Resource (P40 OD028116). We thank the NIH Tetramer Core Facility (contract number 75N93020D00005) for generating the Mafa-A1*063 Gag_386-394_GW9 biotinylated monomers. The following reagent was obtained through the NIH HIV Reagent Program, Division of AIDS, NIAID, NIH: Anti-Simian Immunodeficiency Virus (SIV) gp120 Monoclonal Antibody (B404), ARP-12146, contributed by Dr. Takeo Kuwata. The author(s) utilized the University of Wisconsin – Madison Biotechnology Gene Expression Center (Research Resource Identifier - RRID:SCR_017757) for RNA library preparation and the DNA Sequencing Facility (RRID:SCR_017759) for sequencing.

We are grateful to the WNPRC staff for the exceptional veterinary care provided to the animals throughout this study. The Wisconsin National Primate Research Center is supported by grants P51RR000167 and P51OD011106. This work was funded through the National Institutes of Health [NIH R01 AI108415 and R01 AI124965 (to CVF)] and with federal funds from the National Cancer Institute, National Institutes of Health, under Contract No. 75N91019D00024/HHSN261201500003I. The content of this publication does not necessarily reflect the views or policies of the Department of Health and Human Services, nor does mention of trade names, commercial products, or organizations imply endorsement by the U.S. Government.

