## Supplemental Figures for "CD8+ cells and small viral reservoirs facilitate post-ART control of SIV in Mauritian cynomolgus macaques"

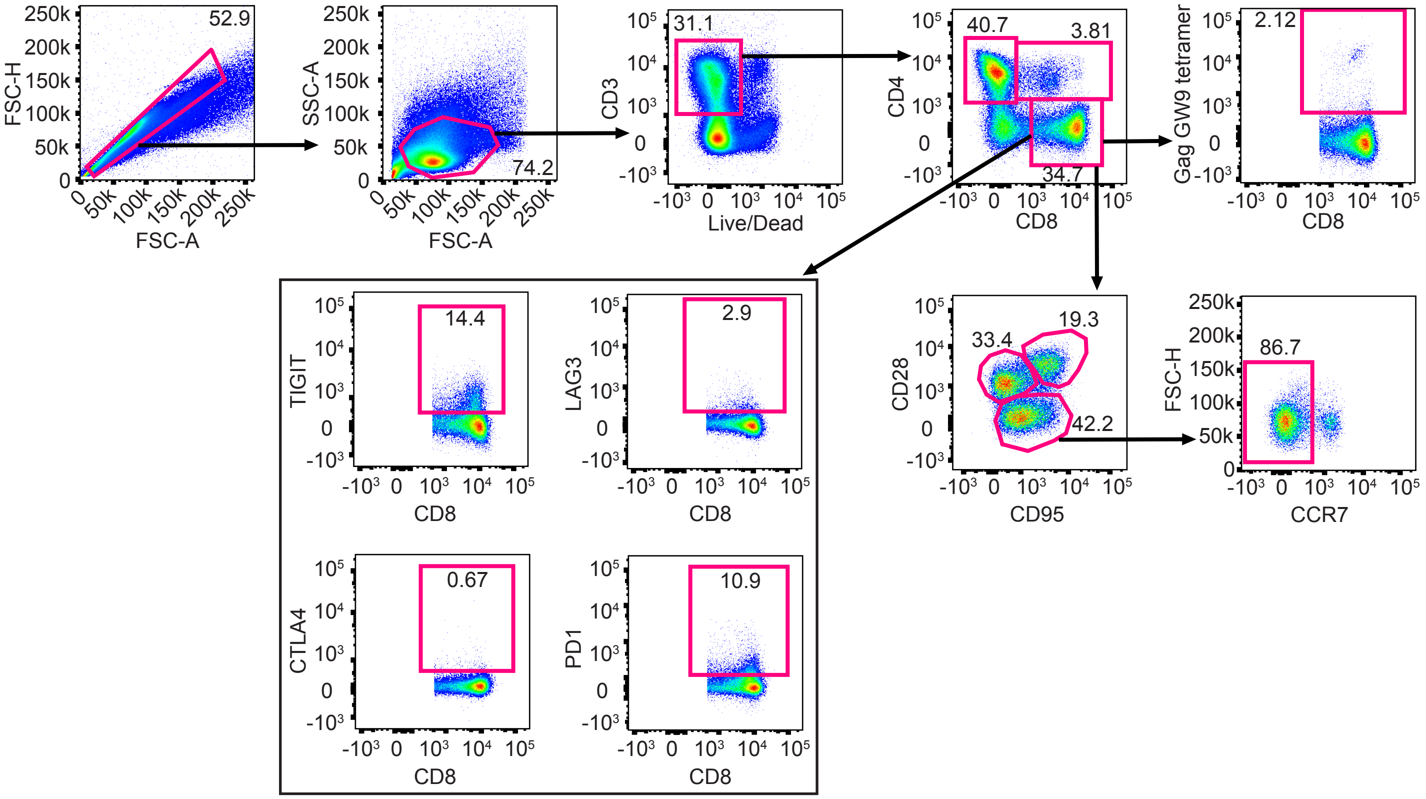


**Supplemental Fig. 1. T cell gating schematic.** Representative gating strategy used to evaluate frequency, memory phenotype, and exhaustion marker expression of bulk CD4+, CD8+, and CD4+CD8+ T cells, and antigen-specific CD8+ T cells.

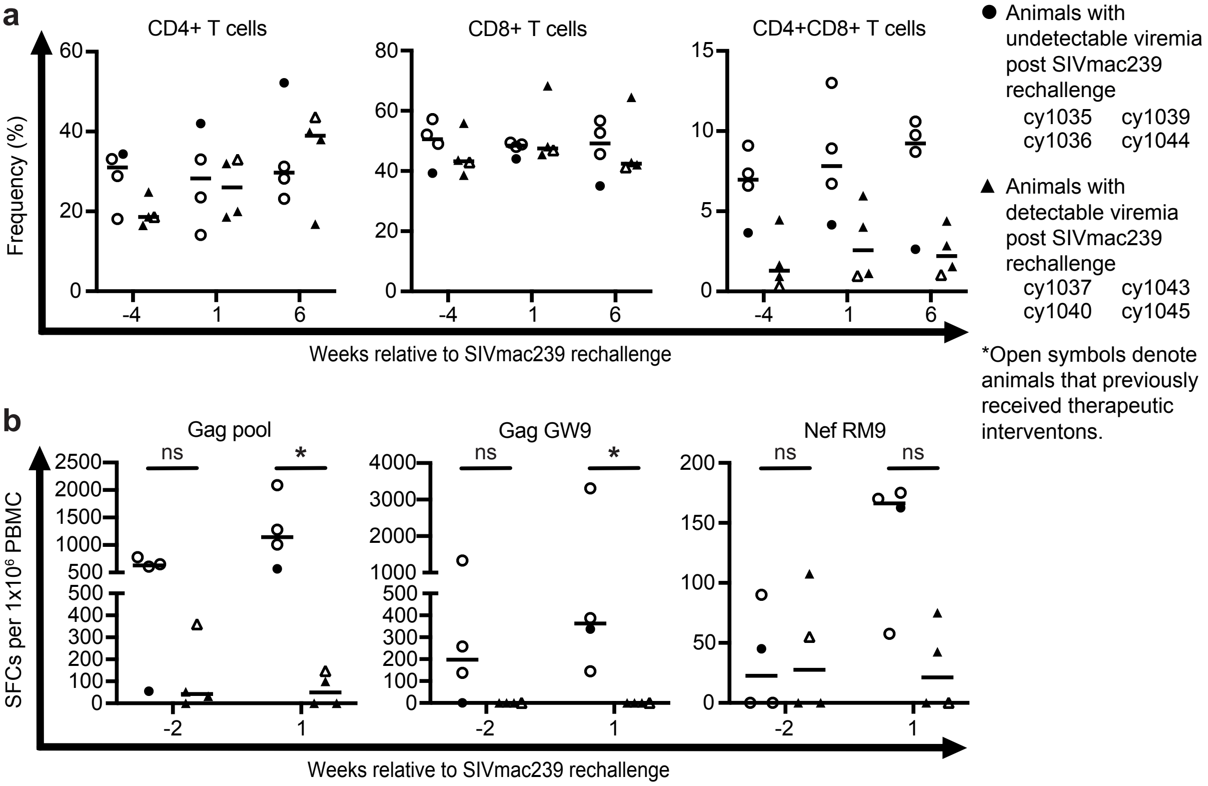


**Supplemental Fig. 2. Immunological changes before and after SIVmac239 rechallenge. a** The frequency of CD4+ T cells (left), CD8+ T cells (center), and CD4+CD8+ T cells (right) before, one week after, and six weeks after SIVmac239 rechallenge in the animals that had undetectable (circles) or detectible (triangles) viremia after rechallenge. Results are displayed for each animal individually with the lines at the median. **b** IFN-γ ELISPOT assays were performed before and one week after SIVmac239 rechallenge to assess responses to $Gag_{386\text{-}394}$GW9, $Nef_{103\text{-}111}$RM9, and a Gag peptide pool in the animals that had undetectable (circles) or detectable (triangles) viremia after rechallenge. Results are displayed for each animal individually with the lines at the median. Animals that previously received therapeutic interventions are displayed as open symbols. * *P*=0.0286. P values were calculated using Mann-Whitney U tests.

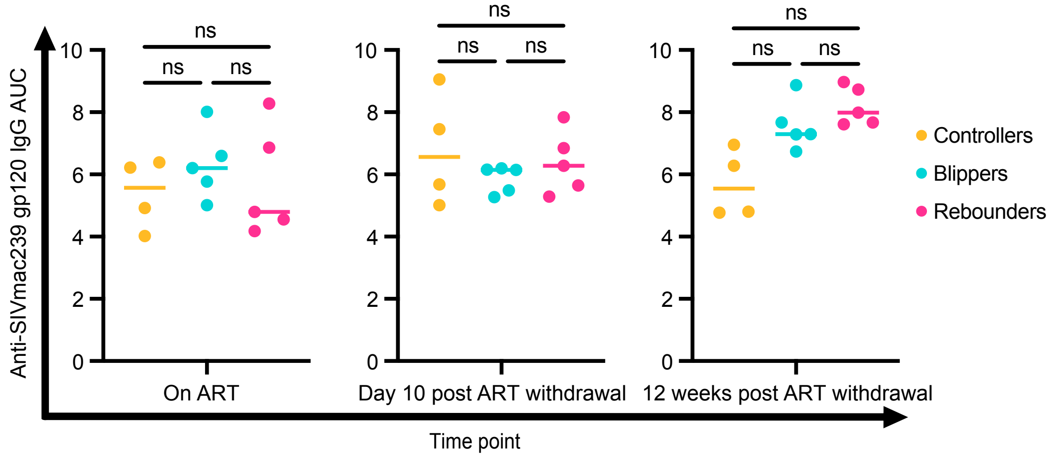


**Supplemental Fig. 3. Anti-SIVmac239 gp120 IgG antibodies from post-ART SIV controllers, blippers, and rebounders.** Area under the curve (AUC) of plasma anti-SIVmac239 gp120 IgG antibodies during ART (left), 10 days after ART withdrawal (center), and 80/81 days after ART withdrawal (right) in the controllers (orange), blippers (teal), and rebounders (pink).

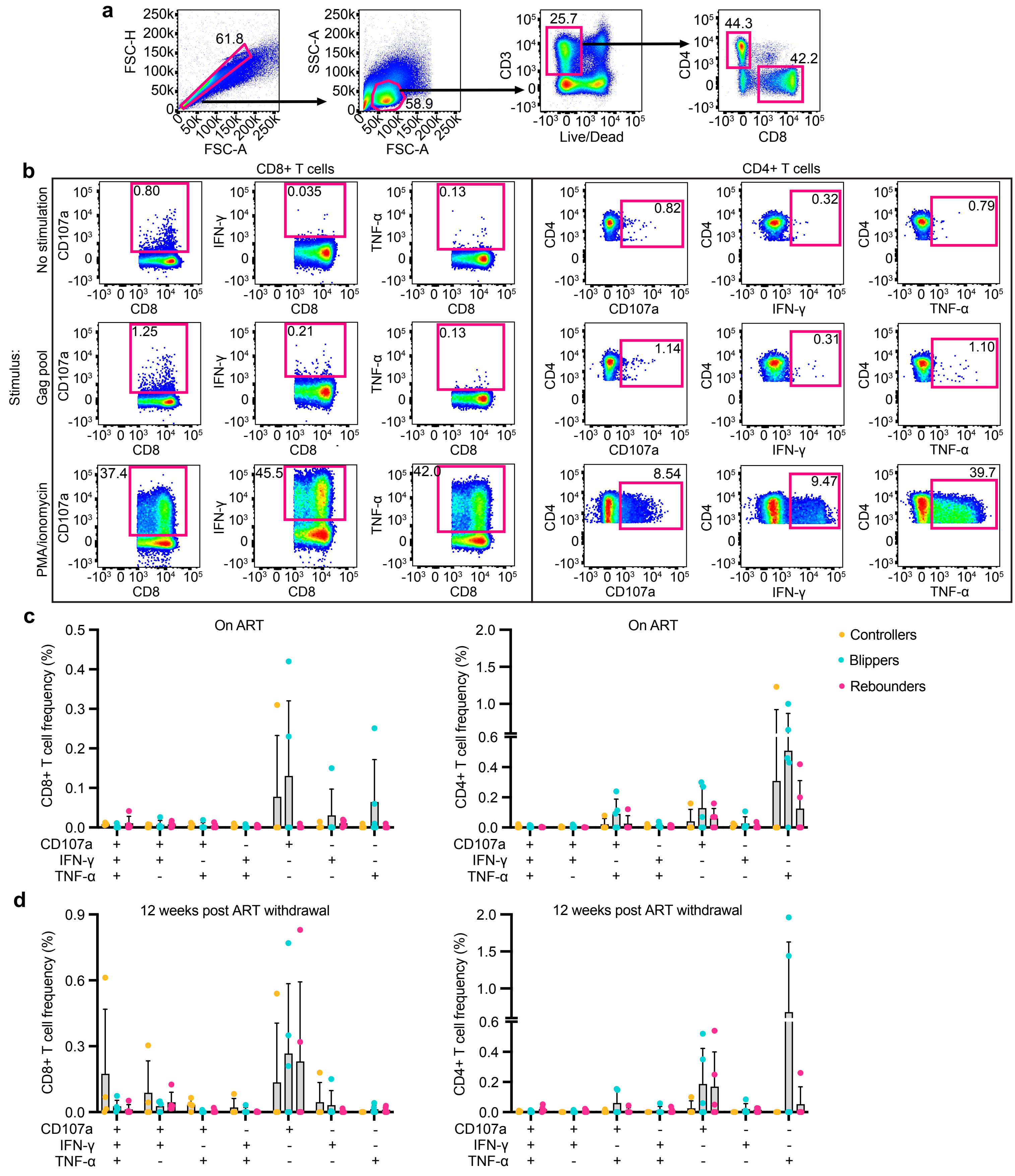


**Supplemental Fig. 4. ICS assay gating schematic and frequencies of polyfunctional CD4+ T cells and CD8+ T cells from post-ART SIV controllers, blippers, and rebounders on ART and after ART withdrawal. a** Representative gating strategy used to distinguish CD4+ and CD8+ T cell populations following overnight incubation with media alone (no stimulation) or a Gag peptide pool. **b** Representative CD107a+, IFN-γ+, and TNF-α+ subpopulations with each respective stimulus are shown for CD8+ T cells (left) and CD4+ T cells (right). **c-d** The frequency of CD8+ T cells (left) and CD4+ T cells (right) from the post-ART SIV controllers (orange), blippers (teal), and rebounders (pink) during ART (**c**) and 12 weeks post ART withdrawal (**d**) expressing CD107a, TNF-α, and/or IFN-γ in response to in vitro Gag stimulation. Responses were calculated with Boolean gates identifying populations expressing permutations of CD107a, TNF-α, and IFN-γ after background subtraction.

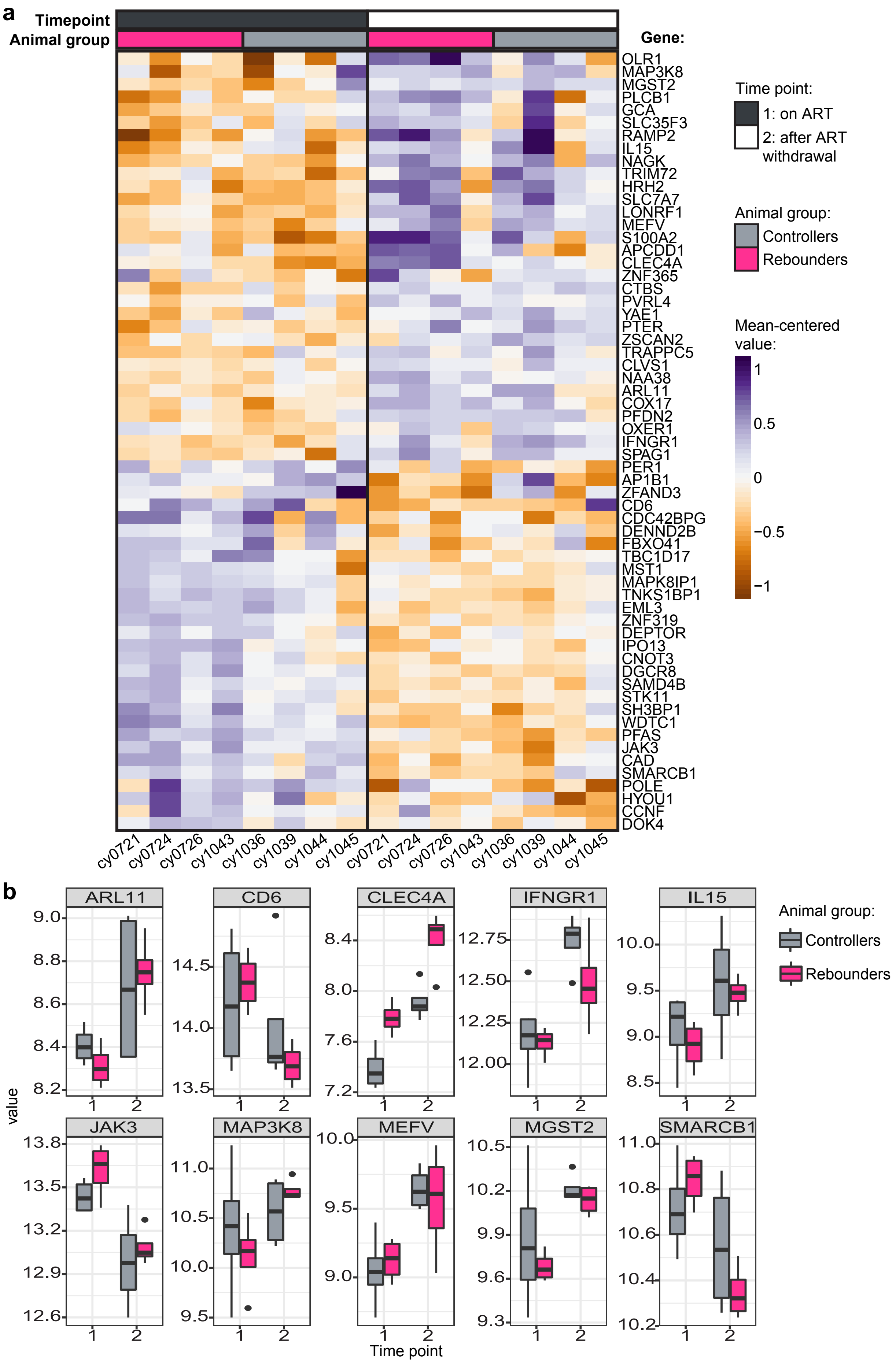


**Supplemental Fig. 5. Differential gene expression of CD8+ T cells from controllers and rebounders on ART and after ART withdrawal. a** Heat map showing the mean-centered value of statistically significant differentially expressed genes between four controllers (gray) and four rebounders (pink) on ART (time point 1, dark gray) and after ART withdrawal (time point 2, white). **b** Box plots showing the values for differentially expressed genes in a that are related to immune function. Genes were considered significant if they had a false discovery rate of < 0.05.

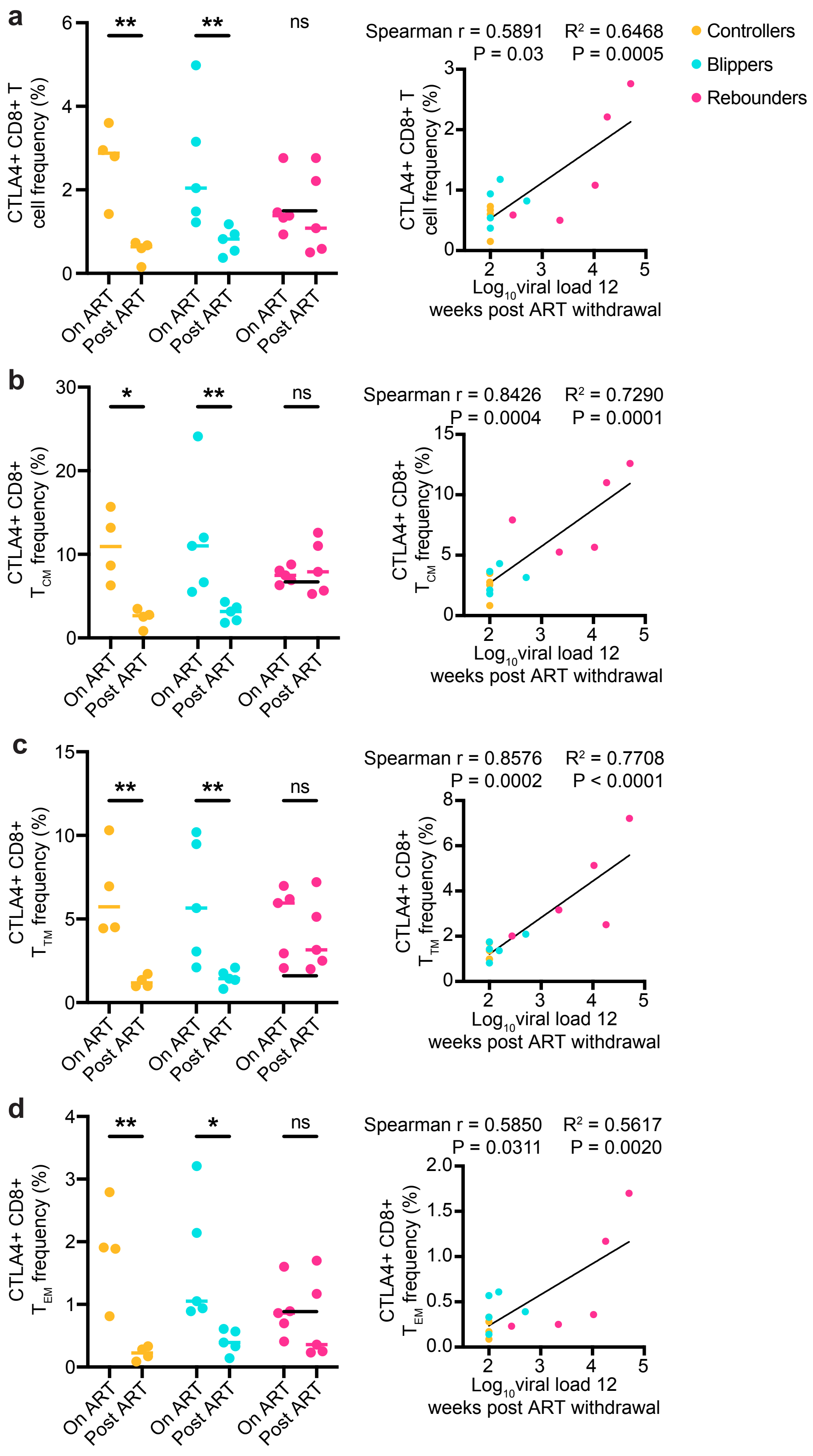


**Supplemental Fig. 6. Decreased CTLA4 expression on CD8+ T cells from post-ART controllers and blippers is associated with lower plasma viral load.** Frequency of **a** bulk, **b** CD8+ $T_{CM}$, **c** CD8+ $T_{TM}$, and **d** CD8+ $T_{EM}$ cells from the post-ART SIV controllers (orange), blippers (teal), and rebounders (pink) during ART treatment (on ART) and 12 weeks post ART withdrawal (post ART). * $P \leq 0.05$, ** $P \leq 0.01$. *P* values were calculated using two-way ANOVA tests with Šídák correction for multiple comparisons. Spearman correlations between the frequencies of each respective CD8+ T cell population and the $log_{10}$viral load 12 weeks after ART withdrawal for all animals are displayed on the right. Spearman r coefficients and *P* values are displayed above each correlation graph. $R^2$ values and best-fit trend lines are also displayed following linear regression analysis.

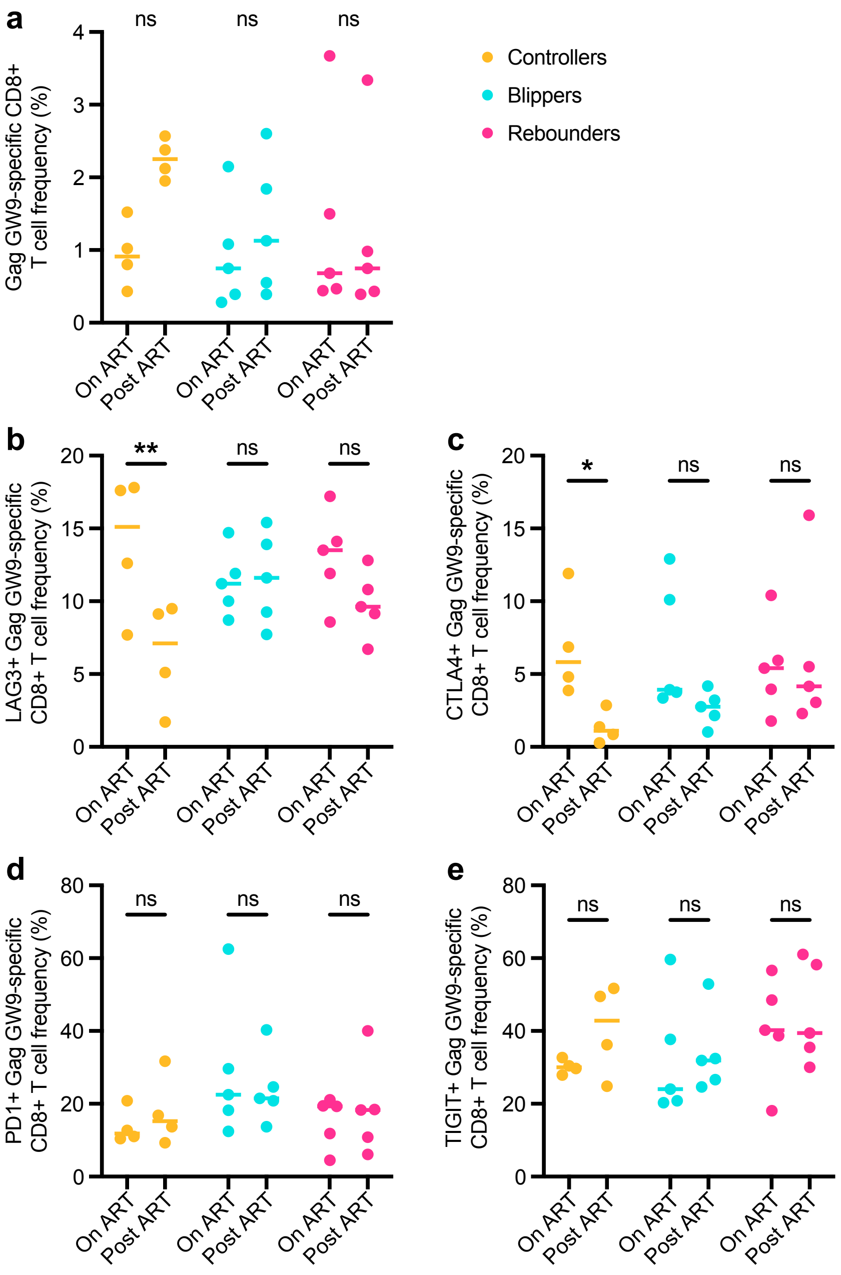


**Supplemental Fig. 7. Frequency and exhaustion marker expression on Gag GW9-specific CD8+ T cells from post-ART SIV controllers, blippers, and rebounders on ART and after ART withdrawal. a** Frequency of Gag GW9-specific CD8+ T cells, and **b-e** Frequency of Gag GW9-specific CD8+ T cells expressing LAG3 (**b**), CTLA4 (**c**), PD1 (**d**), and TIGIT (**e**) in the controllers (orange), blippers (teal) and rebounders (pink) during ART treatment (on ART) and 12 weeks post ART withdrawal (post ART). Results are displayed for each animal individually with the lines at the median. * $P \leq 0.05$, ** $P \leq 0.01$. P values were calculated using two-way ANOVA tests with Šídák correction for multiple comparisons.

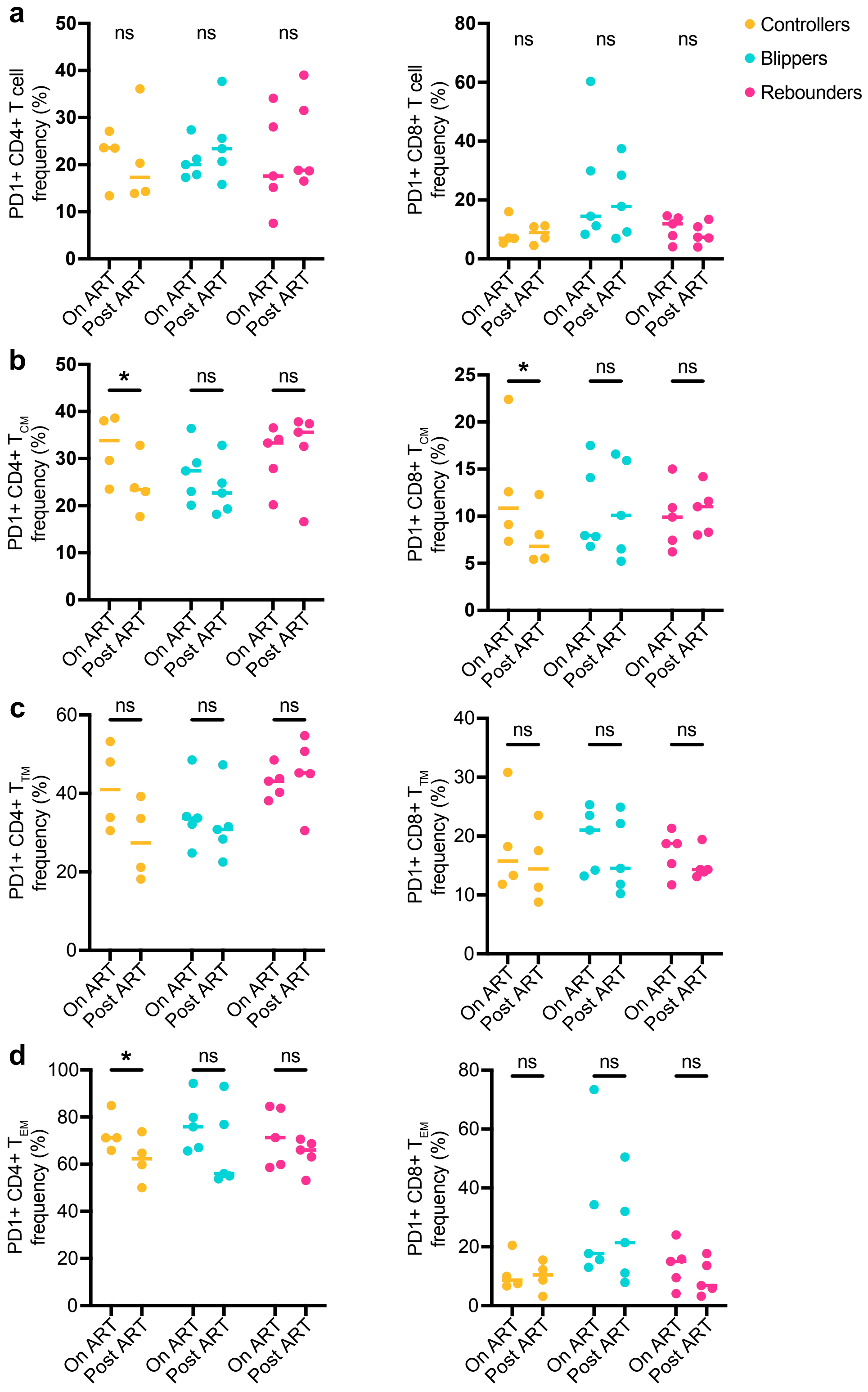


**Supplemental Fig. 8. PD1 expression on bulk, $T_{CM}$, $T_{TM}$, and $T_{EM}$ CD4+ and CD8+ T cells. a-d** The frequency of CD4+ (left) or CD8+ (right) bulk cells (**a**), $T_{CM}$ (**b**), $T_{TM}$ (**c**), and $T_{EM}$ (**d**) expressing PD1 in the controllers (orange), blippers (teal), and rebounders (pink) during ART treatment (on ART) and 12 weeks post ART withdrawal (post ART).
